# Pixel size limit of the PRIMA implants: from humans to rodents and back

**DOI:** 10.1101/2022.06.29.498181

**Authors:** Bing-Yi Wang, Zhijie Charles Chen, Mohajeet Bhuckory, Anna Kochnev Goldstein, Daniel Palanker

## Abstract

**Objective:** Retinal prostheses aim at restoring sight in patients with retinal degeneration by electrically stimulating the inner retinal neurons. Clinical trials with patients blinded by atrophic Age-related Macular Degeneration (AMD) using the PRIMA subretinal implant, a 2×2 mm array of 100μm-wide photovoltaic pixels, have demonstrated a prosthetic visual acuity closely matching the pixel size. Further improvement in resolution requires smaller pixels, which necessitates more intense stimulation.

**Approach:** Here, we examine the lower limit of the pixel size for PRIMA implants by modeling the electric field, leveraging the clinical benchmarks, as well as using a preclinical animal data to assess the stimulation strength and contrast of various patterns. Visually evoked potentials were measured in RCS rats with photovoltaic implants of 100 and 75μm pixels and compared to clinical thresholds with 100 μm pixels. Electrical stimulation model calibrated by these clinical and rodent data was used to predict the performance of the implant with smaller pixels.

**Main Results:** We found that PRIMA implants with 75μm pixels under the maximum safe near-infrared (880nm) illumination of 8 mW/mm^2^ with 30% duty cycle (10ms pulses at 30Hz) should provide a similar perceptual brightness as with 100 μm pixels under 3 mW/mm^2^ irradiance, used in the current clinical trials. Contrast of the Landolt C pattern scaled down to 75μm pixels is also similar under such illumination to that with 100μm pixels in clinical settings, increasing the maximum acuity from 20/420 to 20/300.

**Significance:** Computational model of the photovoltaic subretinal prosthesis defines the minimum pixel size of the PRIMA implants as 75μm. Increasing the implant width from 2 to 3 mm and reducing the pixel size from 100 to 75μm will nearly quadrupole the number of pixels and thereby should significantly improve the visual performance. Smaller pixels of the same bipolar flat geometry would require excessively intense illumination, and therefore a different pixel design should be considered for further improvement in resolution.

## Introduction

Retinal degenerative diseases, such as retinitis pigmentosa (RP) and age-related macular degeneration (AMD), are the major cause of irreversible visual impairment. RP, a relatively rare class of diseases, originates from various genetic disorders, and typically affects patients in their twenties, eventually leading to profound blindness[1]. Atrophic AMD, on the other hand, causes the loss of central vision later in life due to the gradual demise of photoreceptors, a condition called geographic atrophy. This form of advanced AMD affects millions of patients: about 3% of people above the age of 75, and around 25% above 90[2], [3]. Currently, there is no therapy for such scotomata and the loss of sight is permanent.

While photoreceptors degenerate in these diseases, the downstream neurons remain largely intact[4]–[7]. Electrical stimulation of these neurons allows reintroducing information into the visual pathways for restoration of sight. In RP patients, this approach has been demonstrated with an epiretinal prosthesis Argus I/II (Second Sight Medical Products, Inc, Sylmar, CA, USA) [8] and with a subretinal prosthesis Alpha IMS/AMS (Retina Implant AG, Reutlingen, Germany), achieving visual acuity up to 20/546 [9], [10]. In AMD patients, subretinal photovoltaic prosthesis PRIMA (Pixium Vision, Paris, France) recently demonstrated its initial safety and efficacy [11], [12]. The PRIMA implant is a photovoltaic array, where each pixel has two photodiodes connected in series between the active electrode in the center and a return electrode on the circumference (Figure 1A,B). The photovoltaic pixels convert images projected from the augmented-reality glasses into patterns of electric current, stimulating the nearby bipolar cells. The first clinical trial demonstrated a letter acuity closely matching the 100 μm pixel size of the implant (1.17±0.13 pixels), corresponding to the Snellen range of 20/438 – 20/565 [11], [12]. Furthermore, patients simultaneously perceive prosthetic central vision and natural peripheral vision. This inspiring result indicates that smaller pixels may provide higher resolution to benefit a greater number of patients.

**Figure 1.**
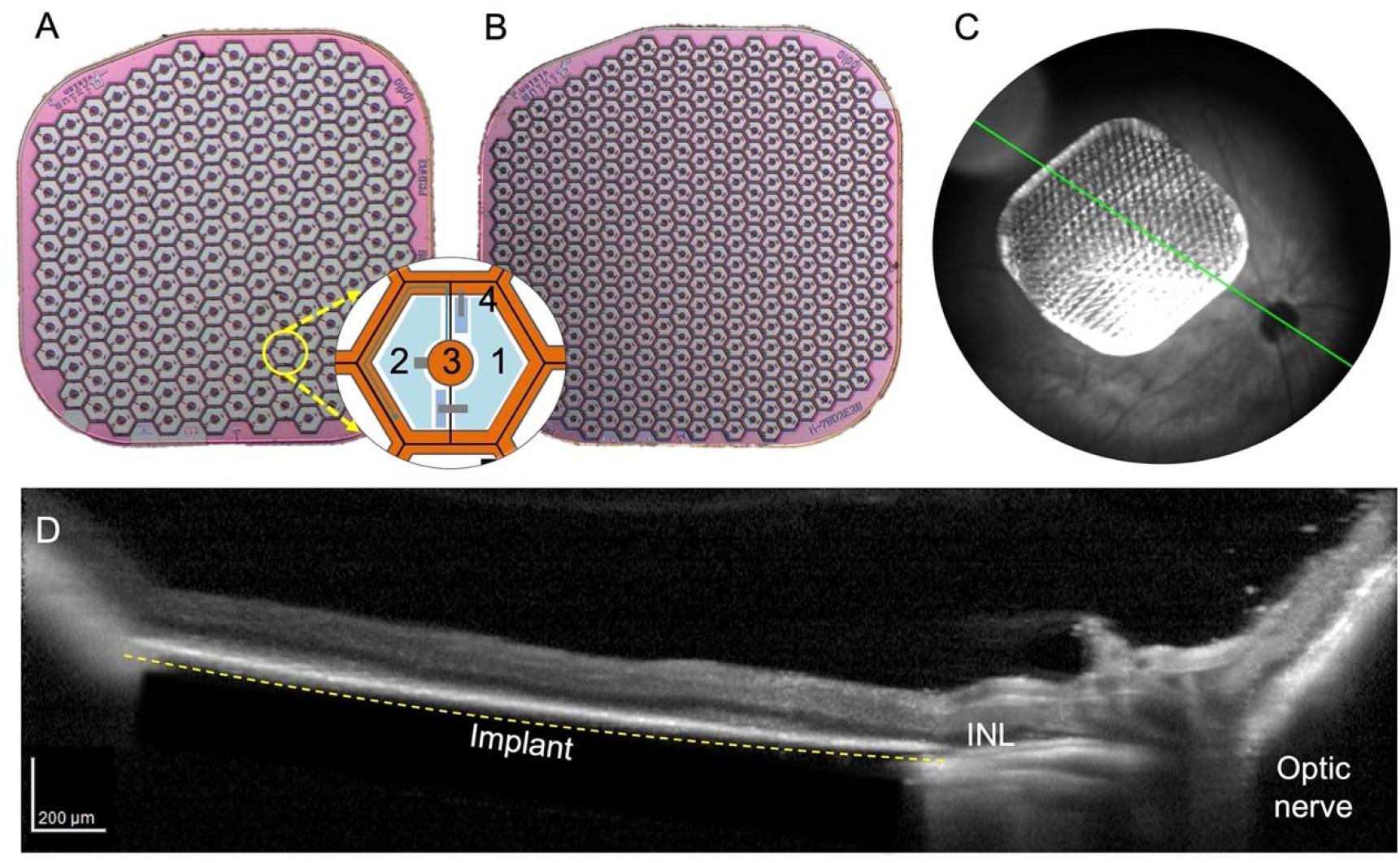
PRIMA photodiode arrays. A) a 100 μm pixel array, B) a 75 μm pixel array. The inset shows the layout of a single pixel, where 1 and 2 are the photosensitive regions of the photodiodes, 3 marks the active electrode, and 4 the local return electrode. C) A fondus image of an implanted 75 μm pixel array in the temporal-dorsal region of the rat retina. D) OCT image of the RCS rat retina with a PRIMA implant several weeks post-op. Dashed line indicates the back side of the implant resting on RPE.

Decreasing the pixel size is not trivial since the retinal stimulation threshold increases nearly quadratically with a decreasing width of bipolar pixels[13], and therefore the lower limit of the pixel size is constrained by the ocular laser safety. For an implant of 3 mm in diameter (the expected dimension in future clinical trials), the steady-state temperature increases by about 2 oC under 5 mW/mm2 NIR illumination[14]. To limit the temperature rise by 1 oC (half of the recommended thermal safety limit of 2°C for active implanted medical devices, according to ISO 14708-1) with a maximum duty cycle of 30% (10 ms pulses at 30 Hz, for instance), the peak NIR irradiance should not exceed 8.25 mW/mm2.

Clinical performance of the 100 mm PRIMA implants demonstrated comfortably bright visual perception under irradiance of 3 mW/mm2 at the maximum pulse duration of 9.8ms and repetition rate of 30 Hz [11], [12]. According to the square scaling of the stimulation threshold with a decreasing size of bipolar pixels in rats[13], pixels of 75 mm would require approximately doubling the light intensity, which should still fit within the ocular safety limit. However, since the human retina is thicker and, unlike the RCS rat retina, the INL is separated from the implant by about 35 mm [11], scaling with the pixel size might be steeper.

In this study, we evaluate the feasibility of the clinical use of 75 mm pixels by relating the implant performance in rats and humans. First, using the electric field model for PRIMA implants, we convert the measured thresholds with 100 μm and 75 μm pixels in rats from the units of irradiance to the voltage step across bipolar cells. Next, using the clinical data with 100 μm pixels and the modeling results, we predict the corresponding thresholds and dynamic range with 75 μm pixels in human. Based on benchmarks derived from the successful 100 μm PRIMA implants and their modeling, this approach allows designing and defining the limits of the next-generation retinal prostheses.

## Methods

### Photovoltaic subretinal prosthesis design

The PRIMA subretinal photovoltaic implants electrically stimulate the second-order retinal neurons, primarily the bipolar cells, to elicit visual perception, thereby enabling restoration of sight[15]. Subretinal photovoltaic pixels activated by light[14], [16] act as an optoelectronic substitute for the lost photoreceptors. To reduce the cross-talk between the neighboring pixels, each one has a local return electrode surrounding the active electrode, as illustrated in Figure 1 A,B. Light-to-current conversion in photoreceptors includes a very strong amplification, up to 6 orders of magnitude: from a few photons per second to pA of current per cell. Since photodiodes do not provide any current amplification and the ambient light intensity is insufficient for electrical stimulation[16], more intense light is required. In addition, capacitive coupling between the electrodes and electrolyte necessitates pulsed charge-balanced stimulation. For these two reasons, the system includes augmented-reality (AR) glasses. Images of the visual scene captured by a video camera are processed and projected by the AR glasses onto a subretinal photodiode array using intense pulsed light. Photovoltaic pixels in the array convert this light into biphasic pulses of electric current, which stimulate the second-order neurons in the inner nuclear layer (INL) – primarily the bipolar cells. To avoid perception of this light by the remaining photoreceptors, a near-infrared (NIR, 880 nm) wavelength is used.

This approach offers multiple advantages: (1) thousands of pixels in the implant can be activated simultaneously and independently; (2) since the pixels are activated by light, no wires are involved, which enables reliable encapsulation of the wireless implant and greatly simplifies the surgical procedure; (3) external camera provides autofocusing and adaptation over a wide range of ambient illumination, as well as adjustable image processing optimized for the dynamic range of the implant; (4) the optical nature of the implant maintains the natural link between the eye movements and image perception; (5) network-mediated retinal stimulation retains many features of the natural signal processing, including the antagonistic center-surround[17], flicker fusion at high frequencies and nonlinear summation of the RGC subunits[14], among others.

### Surgical procedure and animal handling

The 1.5 mm diameter photovoltaic devices were implanted in the subretinal space of the Retinal College of Surgeons (RCS) rats, typically at 6 months of age, when the outer nuclear layer has degenerated completely. The total loss of the outer nuclear layer was confirmed by the optical coherence tomography (OCT; HRA2-Spectralis; Heidelberg Engineering, Heidelberg, Germany) in each animal prior to surgery. The implants were placed in the temporal-dorsal region, approximately 1 mm away from the optic nerve. The fundus image in Figure 1C demonstrates the typical location of an implant, and the OCT in Figure 1D illustrates the retinal reattachment after surgery. A total of 10 animals were implanted with the PRIMA devices consisting of 100 μm (n=5) and 75 μm (n=5) bipolar pixels, as described earlier[18]. Animals were anesthetized with a mixture of ketamine (75 mg/kg) and xylazine (5 mg/kg) injected intraperitoneally. To visualize the retina and the implant, animals were monitored over time using OCT. For measurements of the visually evoked potentials (VEP), each animal was implanted with three transcranial screw electrodes: one electrode over each hemisphere of the visual cortex (4 mm lateral from midline, 6 mm caudal to bregma), and a reference electrode (2 mm right of midline and 2 mm anterior to bregma).

All experimental procedures were conducted in accordance with the Statement for the Use of Animals in Ophthalmic and Vision research of the Association for Research in Vision and Ophthalmology (ARVO) and approved by the Stanford Administrative Panel on Laboratory Animal Care. Royal College of Surgeons (RCS) rats were used as an animal model of the inherited retinal degeneration. Animals are maintained at the Stanford Animal Facility under 12h light/12h dark cycles with food and water ad libitum.

### Electrophysiological measurements

Following anesthesia and pupil dilation, the cornea was covered with a viscoelastic gel and a cover slip to cancel its optical power and maintain good retinal visibility. The implant was illuminated with 880 nm NIR laser (MF-880 nm-400 μm, DILAS, Tucson, AZ) and a digital micromirror display (DMD; DLP Light Commander; LOGIC PD, Carlsbad, CA) was used for a pattern formation. The customized optical system for the pattern projection into the eye was integrated with a slit lamp (Zeiss SL-120; Carl Zeiss, Thornwood, NY) for real-time observation of the retina via a CCD camera (acA1300-60gmNIR; Basler, Ahrensburg, Germany). Light intensity at the cornea was calibrated before and after each measurement session and scaled by the ocular magnification squared to retrieve the irradiance on the implant. Ocular magnification was defined as the ratio between the size of a projected square on the retina and in air.

Visually evoked potentials (VEP) were recorded across the ipsilateral and contralateral visual cortices via the Espion E3 system (Diagnosys LLC, Lowell, MA) at a sampling rate of 2 or 4 kHz, averaged over 250 trials. Corneal signal was simultaneously measured across the ERG electrodes on the cornea and a reference electrode in the nose. The ground electrode was placed in the rat’s tail. The corneal signal served as a template for stimulus artifact removal in the VEP waveforms. The noise floor was extracted from the VEP measurements without illumination. VEP amplitude was measured as the peak-to-peak value in the 150 ms window after the stimuli onset (Figure 2A). Stimulation thresholds were measured with full-field and partial-field (0.65 × 0.65 mm) illumination (Figure 3C) with 10 ms pulses at 2 Hz repetition rate and irradiances ranging from 0.024 to 4.7 mW/mm^2^ on the retina. To determine the strength-duration relationship of the stimulation threshold, we found the minimum stimulation strength necessary to elicit a significant VEP response at pulse durations ranging from 0.5 to 20 ms.

**Figure 2.**
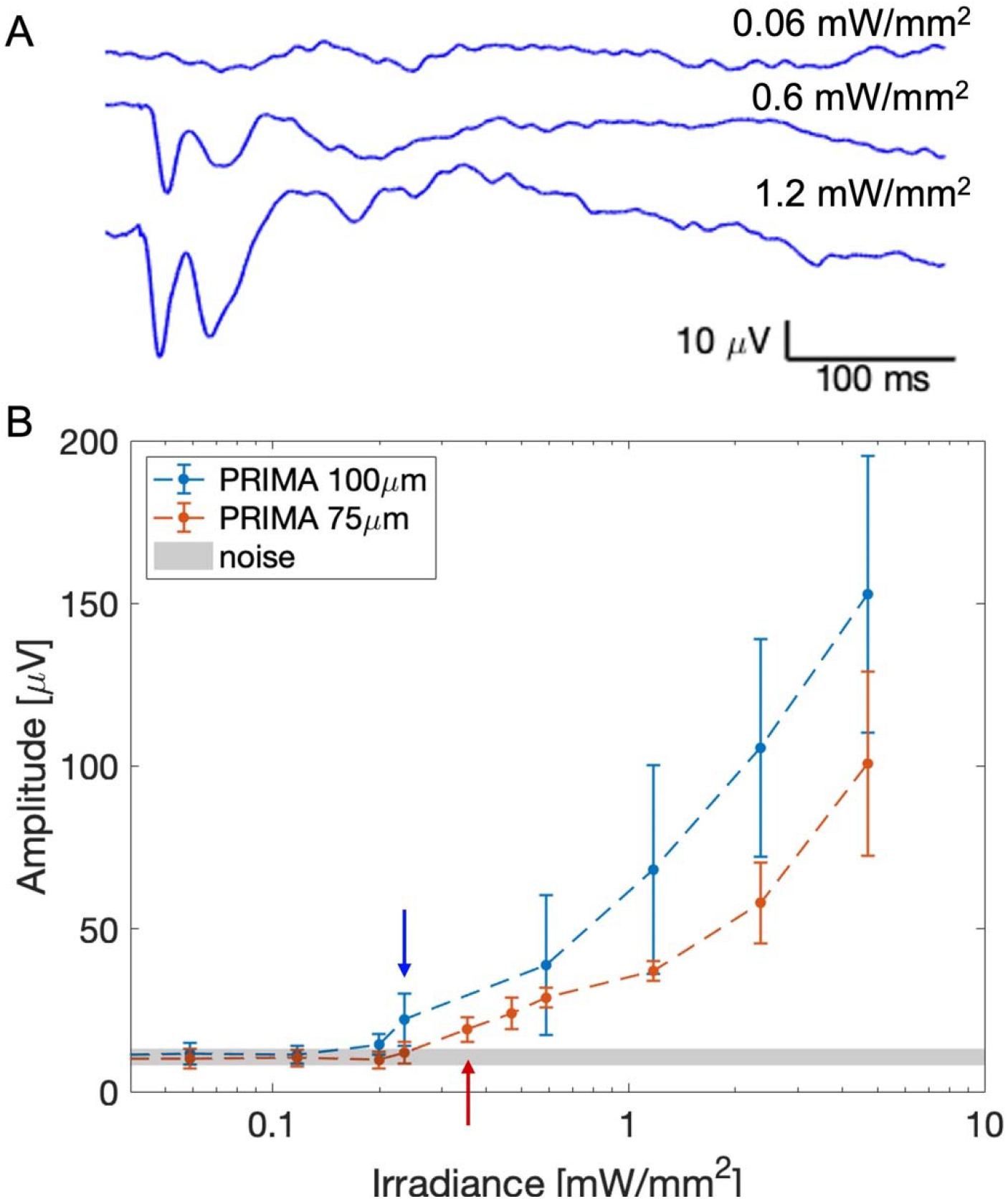
A) Example VEP waveforms measured in an RCS rat with a 75 μm implant. The top waveform is no different from noise (measured with no NIR stimulus), while the middle and bottom traces demonstrate above-threshold waveforms. B) The peak-to-peak VEP amplitude as a function of irradiance. The arrows point to the first irradiance level yielding an amplitude above the noise band for a given pixel size.

**Figure 3.**
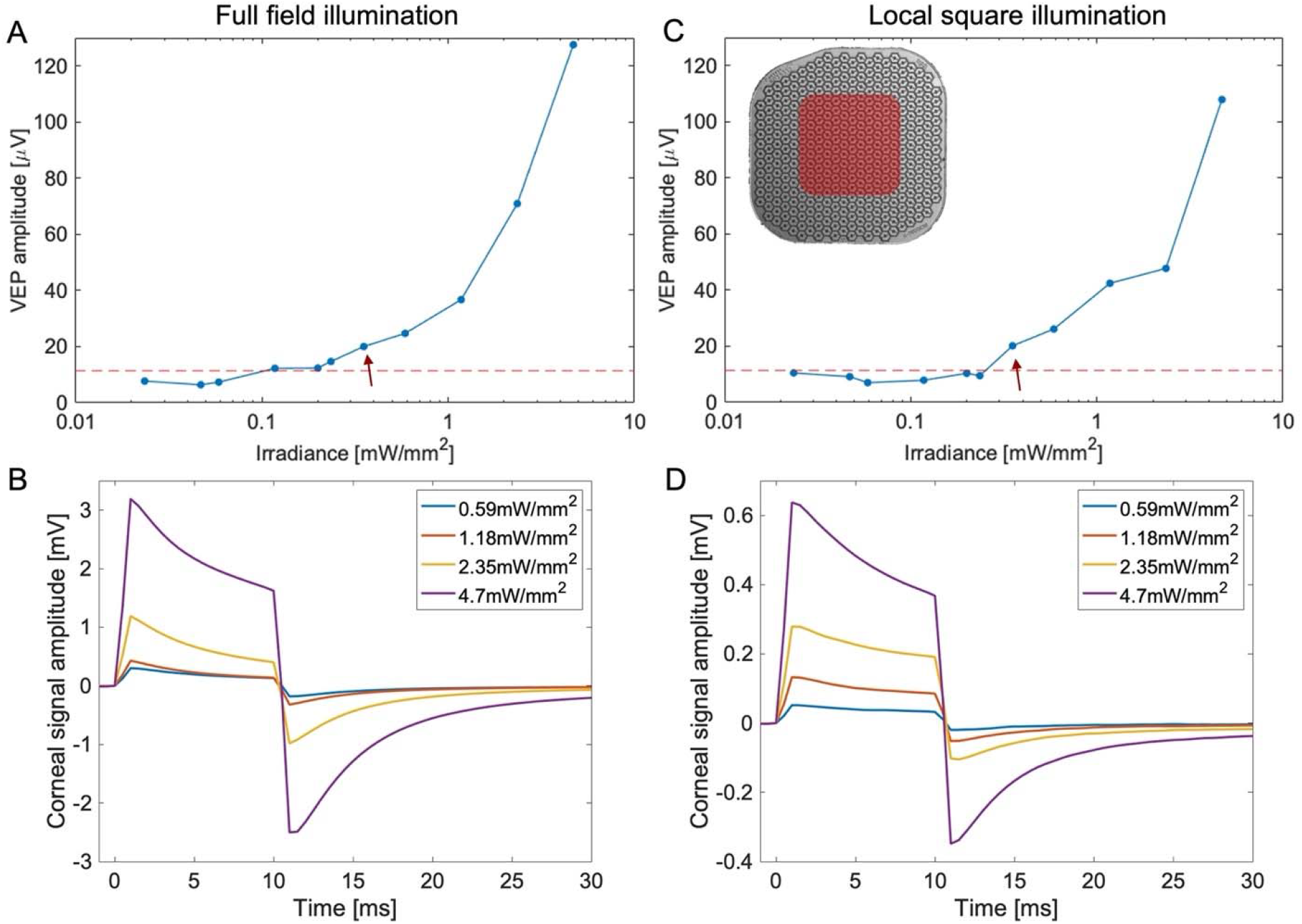
Example of the VEP amplitude and corneal signal measured with full-field illumination (A-B) and with a 0.65 × 0.65mm square illumination (C-D).

### Electric field modeling

To relate the optical stimuli to the electric potential across bipolar cells in the network-mediated retinal stimulation[19], we represented the electric field in the retina by the weighted sum of the pre-computed basis of elementary fields, each of which corresponds to one electrode injecting a unitary current[20]. The weights are the current injections of the pixels, calculated from the light pattern on the implant as a function of time, with RPSim, a circuit simulator based on Xyce[21]. More details regarding the computational framework for assessing the dynamics of the photovoltaic implants[22] are described in the companion paper[23]. We assumed the average length of the bipolar cells to be equal to the distance between the bottom of the INL and the middle of the inner plexiform layer (IPL), which was about 52 μm, as measured by OCT.

## Results

### Electrophysiological performance of PRIMA implants in rodents

To compare the implants’ performance in humans and rodents, we examined the electrophysiological response in rats elicited by 100 μm PRIMA devices. To better fit the size of the RCS rats eye, implants were 1.5 mm in width instead of the 2 mm ones used in clinical trials[11], [12]. Similar devices with 75 μm pixels (Figure 1 A,B) were also implanted and evaluated in-vivo. Figure 2A showcases some typical VEP waveforms measured in RCS rats with 75 μm pixels at various light intensities. Below the stimulation threshold, the waveforms appear indistinguishable from noise. As NIR (880nm) light intensity increases, the signature peaks start appearing and grow in amplitude, while retaining the timing. To determine the stimulation threshold, we swept across a wide range of irradiances with 10 ms pulses at 2 Hz repetition rate. Figure 2B summarizes the peak-to-peak VEP amplitude as a function of irradiance on the retina. The average stimulation threshold was 0.23 +/-0.02 mW/mm^2^ with 100 μm pixels, and 0.38 +/-0.06 mW/mm^2^ with 75 μm pixels. These are conservative estimates, as we take the first points where VEP amplitudes rise above the noise floor, without extrapolation. Note that the VEP amplitudes at the highest irradiance (5 mW/mm^2^)[12] with 75 μm pixels are similar to those measured at half that irradiance (2.5 mW/mm^2^) with 100 μm pixels. This indicates that perceptual brightness with 75 μm pixels under twice higher irradiance might be similar to that with 100 μm pixels, which in the current clinical trials produce comfortably bright percepts with irradiance of about 3 mW/mm^2^[11], [12].

The perceptual threshold in human patients was measured using a 16-pixel-wide circular spot (1.6 mm in diameter), with 9.8 ms pulses at 10 Hz repetition rate [12], [24]. In rodents, we apply similar pulse duration and repetition rate, but using full-field illumination. To verify whether full-field and partial-field illumination provide similar thresholds, we measured the VEP also using a local square illumination of 0.65 × 0.65 mm, as illustrated in Figure 3C. Despite the lower corneal signals with a partial-field illumination (Figure 3D,B), the cortical response amplitude was found to be similar (Figure 3 A,C). This indicates some additional signal amplification in the brain, which is not as sensitive to the size of large stimuli on the retina. Therefore, electrophysiological stimulation thresholds derived from full-field illumination may be used for comparison with the perceptual thresholds detected with a large spot illumination.

For further comparison with the clinical results, we also characterized the strength-duration (S-D) relationship of the stimulation threshold with 75 and 100 μm pixels in rats. Figure 4A illustrates the stimulation thresholds in terms of light intensity as a function of pulse duration, fitted with the Weiss function for 100 μm and 75 μm implants. The resulting chronaxie for 100 μm pixels was 2.3 ms, in excellent agreement with the chronaxies measured in the PRIMA clinical trial, two examples of which are shown in Figure 5A. Rheobase in the rat measurements (0.17 mW/mm^2^), however, was significantly lower than even in the best case of human patients (0.4 mW/mm^2^). This is likely due to the fact that, unlike in RCS rats, INL in human patients is separated from the implant by about 35 μm of the debris[12].

**Figure 4.**
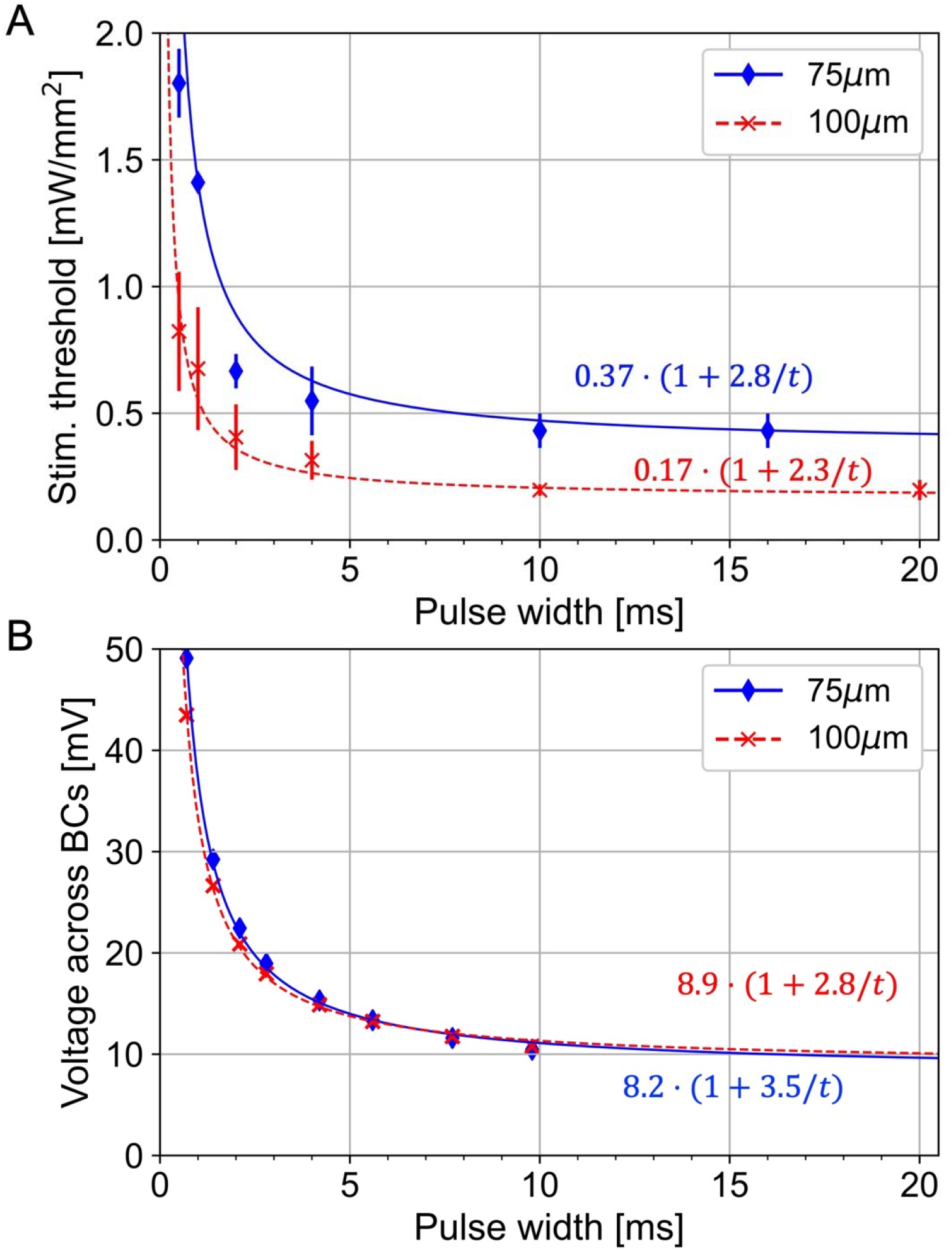
A) The irradiance thresholds as a function of pulse duration measured in RCS rats with 100 μm and 75 μm pixel arrays. The solid and dashed lines show the fitted strength-duration curves according to Weiss equation. B) The converted strength-duration curves in units of voltage across bipolar cells in rat retina.

**Figure 5.**
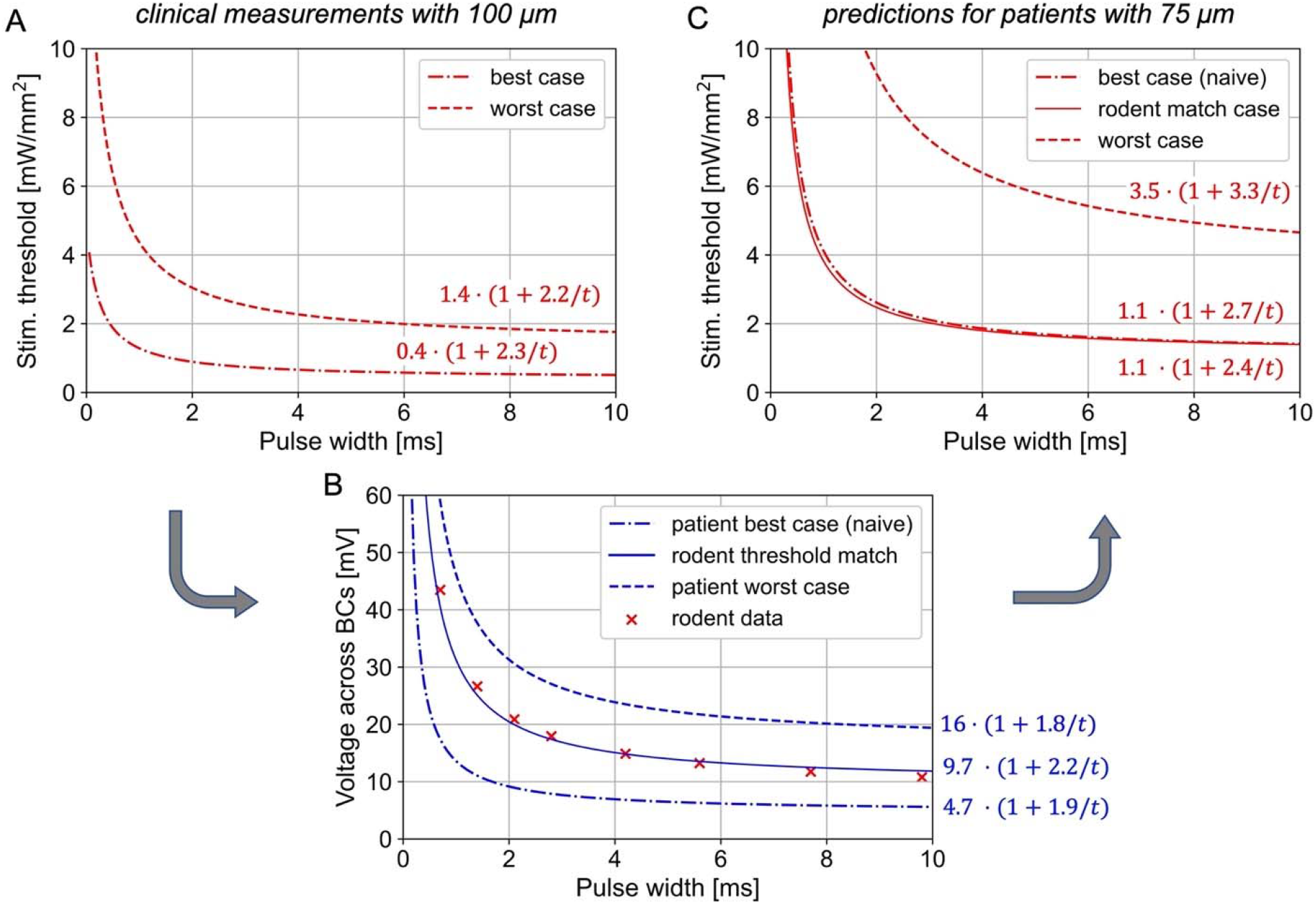
Strength-duration (S-D) relationships for human thresholds with 100 and 75 μm pixel arrays. A) Perceptual thresholds for the best and worst cases measured with 100 μm PRIMA implants. B) Stimulation thresholds in terms of a voltage drop across bipolar cells (BCs) for the best and the worst case, converted from A. The middle line represents the best-case thresholds with 100 μm implants adjusted to 20 μm separation between the BCs and the implant best case to match the rodent data (from Figure 4B). C) The predicted S-D curves for 75 μm pixels in all three cases, converted from the voltage to irradiance.

### Predictions of clinical performance with 75 μm PRIMA implants

Bipolar cells are the target neurons in our subretinal approach, and we have shown previously that depolarization of their axonal terminals can be estimated from the voltage step between their top and bottom boundaries[19]. To assess this voltage step, we converted the stimulation thresholds from the irradiance to voltage using the model of electric field generated by photovoltaic arrays in electrolyte[23]. Figure 4B shows the S-D curves for 100 μm and 75 μm pixels measured in RCS rats, converted into the voltage across BCs, estimated to extend from the bottom of INL (starting about 5μm above the implant) to the middle of IPL, at 57 μm height. Despite the very different levels of light intensity shown in Figure 4A, the voltage S-D curves are nearly identical, confirming that the same voltage step is required for BC stimulation, regardless of the implant configuration. The rheobases for these two fits are in the range of 8-9 mV, and with 10ms pulses, the threshold voltage is about 11 mV.

To predict the clinical performance of PRIMA implants with 75μm pixels, we first convert the human stimulation thresholds with 100 μm pixels from light intensity into the voltage step across bipolar cells, and then calculate the light intensity required to generate the same voltage with 75μm pixels in human retina. Figure 5A shows the lowest and the highest stimulation thresholds in AMD patients reported to date with the 100 μm PRIMA implants [12], which we use as the “best” and “worst” cases to estimate the range of responses with future implants. The chronaxie of the S-D relationships are very similar between these two boundary cases (2.2 and 2.3 ms) while the rheobase varies very significantly: by a factor of 3.5. Conversion to the corresponding voltage thresholds in human retina, where we assume the INL to begin 35 μm above the implant and the middle of IPL at 87 μm, based on clinical OCT images [12], is shown in Figure 5B. For the worst case, a rheobase is about 16.4 mV, roughly twice the rheobase measured in rodents, which may be due to the reduced neural excitability in the badly degenerated retina. In the best case, the stimulation thresholds are more than three times lower, with a corresponding rheobase of 4.7 mV. This value is below the approximate turn-On voltage of the Ca ion channels (> 8 mV)[25], [26]. This discrepancy could be reconciled if we assume that dendrites of the bipolar cells in the best-case patient extend below the cell bodies, thereby reducing the distance from the implant. If we assume that the BC stimulation threshold in human retina in the best-case scenario is similar to that in rats (and matches the expected turn-On voltage of Can channels), the distance between the dendrites and the implant surface should be about 20 μm. This S-D curve, also shown in Figure 5B, lies in between the best and the worst cases, with a rheobase of 9.7mV.

Given the voltage step across bipolar cells, we can now calculate the light intensity required for generating such a voltage drop with 75 μm pixel arrays in human retina. Figure 5C shows the predicted stimulation thresholds in units of irradiance as a function of pulse duration for the two boundary cases. For the worst-case scenario, the rheobase is 3.5 mW/mm^2^ – 2.5 times higher than in the worst case with 100 μm pixels. This increase closely matches the 2.59 ratio of the photosensitive area in 100 and 75 μm PRIMA pixels: 4075 μm^2^ vs. 1576 μm^2^, respectively. This similarity indicates that the same current per pixel is required to produce the same voltage drop across the BCs with these two arrays. This, in turn, indicates that higher current density per unit area of the implant with smaller pixels is compensated by stronger field confinement with denser return electrodes grid. With the irradiance of 8 mW/mm^2^, stimulation should exceed the worst-case threshold with pulses longer than 2.6 ms, which is similar to the minimum pulse duration with 100 μm pixels under 3 mW/mm2 used in the clinical trial [12].

For the best-case scenario, required irradiance can be predicted two ways. First, using a ‘naive’ voltage threshold, which assumes the BCs residing 35 μm above the implant and having the rheobase of 4.7 mV. The more physiologically realistic voltage thresholds, as described above, correspond to BCs extending their dendrites down to 20 μm above the implant and having the rheobase of 9.7mV. Interestingly, both models yield very similar S-D curves in units of light intensity, as shown in Figure 5C. Here again, the ratio of the average rheobase with 75μm pixels (1.05 mW/mm^2^) to that with 100 μm pixels shown in Figure 5A (0.4 mW/mm^2^) is about 2.6, like in the worst-case scenario. Therefore, clinical performance of the 75 μm implants in the best and the worst cases under 8 mW/mm^2^ irradiance should be similar to that in the clinical trial with 100 μm pixels, where irradiance was about 2.6 times lower – 3 mW/mm^2^[11], [12].

Prosthetic visual acuity in the clinical trials was assessed using the Landolt C optotype, projected with 9.8ms pulses at 30 Hz. Theoretically, the minimum resolvable font size corresponds to the gap of 1 pixel in the letter C. Patients resolved the fonts in the range of 1.04 – 1.3 pixels per gap, with the average of 1.17 pixels [12]. To check the expected resolution with 75 μm pixels, we calculated the electric potential across bipolar cells when a Landolt C font with a gap of 1.2 pixel (90 μm) was projected onto the photovoltaic array with the irradiance of 8 mW/mm^2^ and 9.8 ms pulses at 30 Hz (Figure 6). As with the stimulation thresholds, we calculated the field maps for two scenarios: if the BCs are separated from the implant by 35 μm and by 20 μm. With the 35 μm separation, we outlined two thresholds for 10 ms pulses: for the best (5.6 mV) and the worst (18.9 mV) cases. For the 20 μm separation, assumed only for the best case, the threshold is 11.8 mV. As shown in Figure 6B and C, the predicted contours of visibility for the best case in two scenarios are very similar. The worst case, as shown in Figure 6B, represents a much narrower letter with a wavy outline. However, with the eye movements, this wavy structure may average out and hence not be noticeable by the patients. Amplitude of the electric potential in these maps (57 mV for 20 μm and 28 mV for 35 μm height) is very close to that calculated for the 100 μm pixels under 3 mW/mm^2^ irradiance: 62 mV and 31 mV, respectively[23]. These findings indicate that with 75 μm pixels under 8 mW/mm^2^ irradiance, the 1.2 pixel gap in Landolt C should be resolvable, with a similar perceptual brightness of the pattern as in the current clinical trials.

**Figure 6.**
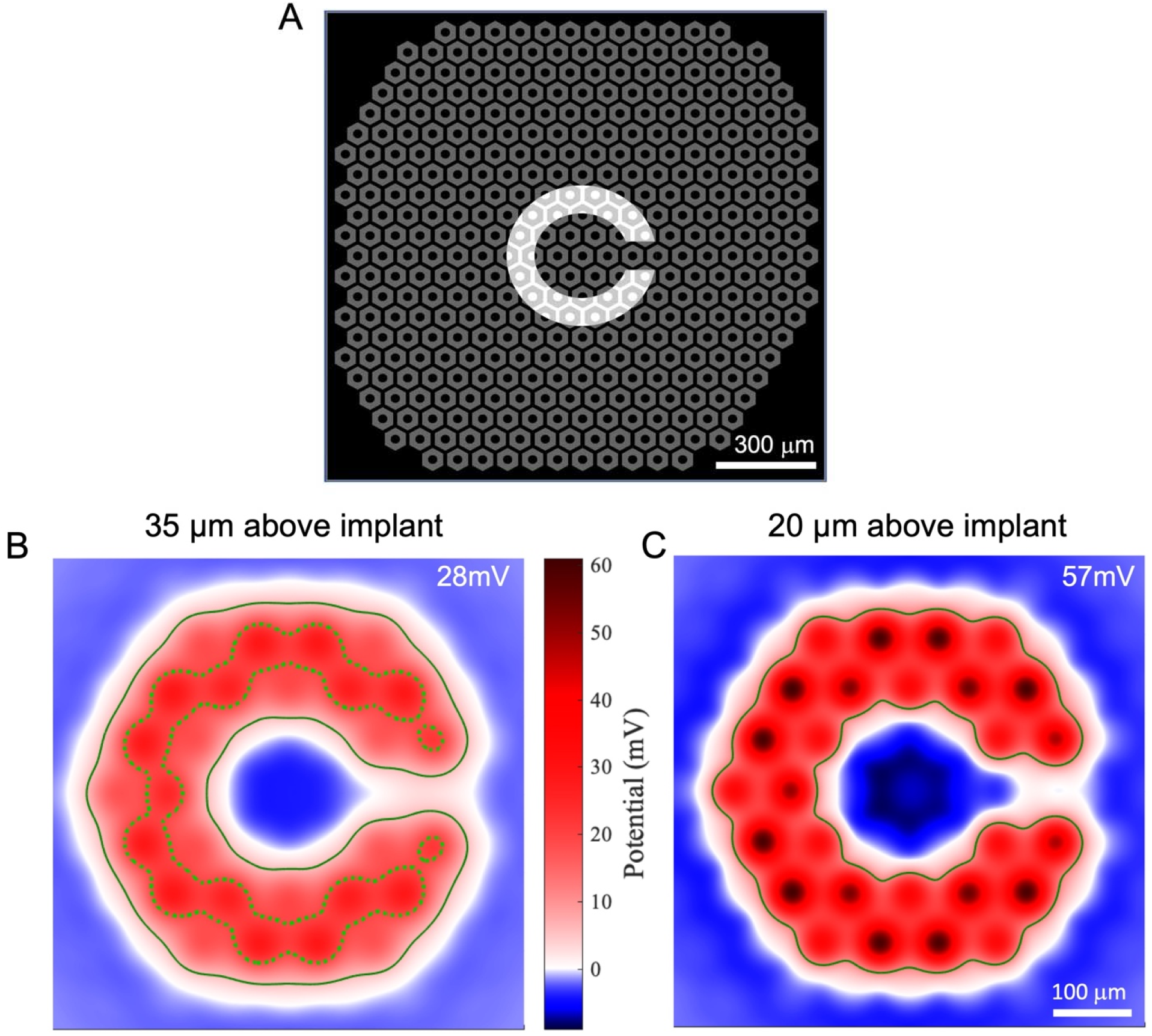
Landolt C pattern stimulation with 75 μm pixels. A) A Landolt C pattern projected on top of a 75 μm pixel array. The width of the gap in C is 90 μm – 1.2 times the pixel width. B) The simulated field map for BCs separated by 35 μm from the implant, whose terminals end in the middle of the inner plexiform layer (87 μm above implant surface). The solid line marks the base-case threshold (5.6 mV), while the dotted line, the worst-case threshold (18.9 mV). The label 28 mV indicates the highest voltage step in this map. C) The field map for BCs separated from the implant by 20 μm, while the location of axonal terminals remain the same as in B. The solid line marks the threshold (11.8 mV) for the best-case at this height. The label 57 mV indicates the highest voltage step here.

## Discussion

The predicted stimulation thresholds with 75 μm bipolar pixels are higher than those measured with 100 μm pixels by a factor of about 2.5, approximately matching the ratio of the photosensitive areas in these pixels. This means that the same current per pixel is required for stimulation, as would be with the electric field of a point source. Such approximation is expected with small electrodes and cells located closer to the active electrode than the distance to return electrode, which is the case with RCS rat retina, where bipolar cells are within a few micrometers from the implant. In AMD patients, however, INL is separated from the implant by about 35 μm[12]. With a pattern activation (Landolt C or a large spot) at distances comparable to or exceeding the pixel radius, higher current density per unit area generated with smaller pixels should increase the electric potential, but at the same time, the electric field is diminished by higher density of the return electrodes. These two effects counteract each other, resulting in the threshold scaling still according to the constant electric current per pixel.

Perceptual brightness and contrast of the patterns with 75 μm pixels at 8 mW/mm^2^ is expected to be similar to that with 100 μm pixels at 3 mW/mm^2^ irradiance in humans. Since 8 mW/mm^2^ with a 30% duty cycle is close to the NIR retinal safety limit, further reduction in pixel size of the same geometry is unlikely. In rodents, where INL is closer to the implant, we demonstrated a prosthetic visual acuity matching the pixel pitch even with 55 μm bipolar pixels[27], but for human retina, achieving similar efficacy with 55 μm pixels would require light intensity exceeding the safety limits.

Bipolar pixels effectively prevent the cross-talk between neighboring pixels, but they do it by overly constraining the electric field. Placing a local return electrode inside each pixel creates a dark ring between the adjacent bright spots in the pattern. Ideally, the electric field should be confined such that the bright pixel creates a stimulation spot, while the dark pixel does not, without a dark boundary around the active pixels. This can be achieved using current steering, if the active electrodes on monopolar pixels could be used for both injecting and collecting the current, i.e. as the active and return electrodes[18], [28]. Additionally, 3-dimensional electrodes can improve the electric field penetration into the tissue, thereby maintaining a low stimulation threshold and good contrast, such as the honeycomb and pillar layouts[13], [29].

## Conclusions

Under the maximum safe NIR irradiance (8mW/mm^2^ with 30% duty cycle), the 75 μm PRIMA implants should provide a similar perceptual brightness to that with the 100 μm implants under 3 mW/mm^2^, which is used in the first clinical trials. It should also maintain sufficient contrast for resolving a Landolt C with a gap width of 1.2 pixel, corresponding to the visual acuity of 20/380. A decrease in the pixel size from 100 to 75 μm and an increase in the implant width from 2 to 3 mm will nearly quadruple the number of pixels in an implant, which should significantly improve its visual performance. Further decrease of the pixel size in flat bipolar geometry would require exceedingly bright illumination, and therefore a different electrode geometry should be considered for further improvement in resolution.

## Acknowledgements

The authors wish to thank Pixium Vision for providing the 100 μm and 75 μm implants. The financial support for this research was offered by National Institutes of Health (Grants R01-EY-027786 and P30-EY-026877), the Department of Defense (Grant W81XWH-19-1-0738), AFOSR (Grant FA9550-19-1-0402), Wu Tsai Institute of Neurosciences at Stanford, and an unrestricted grant from Research to Prevent Blindness.

